# Assessing spatial segregation of beluga whales (*Delphinapterus leucas*) in Western Hudson Bay estuaries

**DOI:** 10.1101/2021.08.05.455325

**Authors:** Kristin Westdal, Jeremy Davies, Steve Ferguson

## Abstract

Segregation of older adult males from females and immature males is known to occur in some beluga whale populations, but it is unclear if adults accompanied by calves segregate in Hudson Bay, where the largest summering population is found. Using imagery from a photographic aerial survey conducted in August 2015, we considered a number of environmental variables that might explain distribution by age class of beluga near two of three main estuaries (Churchill and Seal River) used by Western Hudson Bay belugas in the summer season. Belugas were identified and classified by age manually using an identification decision tree and GPS coordinates were plotted in ArcGIS. Distribution by age class was examined in relation to distance to coastal habitat and bathymetry to test the predation risk hypothesis, sea surface temperature (thermal advantage hypothesis), and extent of river plume (forge-selection hypothesis). Habitat characteristics and the proportion of age classes in both estuaries were similar between age class groups (with and without calves) indicating no segregation and suggesting the environmental data assessed were not driving patterns of distribution and density of age classes at the spatial and temporal scale being investigated. Results provide a greater understanding of spatial patterns of beluga whale habitat use in western Hudson Bay and information useful in conservation and management advice.

## Introduction

Social and spatial segregation of sexes is common amongst vertebrates that live in groups [1].Whale species such as the North Atlantic right whale (*Eubalaena glacialis*), North Pacific gray whale (*Eschrichtius robustus*) and bowhead whale (*Balaena mysticetus*) are known to have nursery areas where females feed and take care of their young [2, 3, 4]. The right whale is thought to prefer shallow coastal areas, the gray whale coastal lagoons, and the bowhead is known to use a separate shallow-water part of their summer range [2,4]. Although segregation of older adult males from immature males and females is known to occur in beluga whales (*Delphinapterus leucas*) in the western Arctic [5], and habitat preferences differ for males and females in the central Arctic [6] it is unclear if adult accompanied by calves segregate for periods of time during the year in the eastern Arctic. In Hudson Bay, beluga family groups have been shown to migrate together and females were found to be more likely to travel in a family group than males [7].

Western Hudson Bay belugas inhabit offshore areas with dense pack ice in winter and prefer shallow warm water estuaries in the summer season undertaking migrations of more than 2,000 km round trip between their summer and winter habitat, similar to long distance migrations of other beluga populations [8]. Although somewhat debated in the literature, long held hypotheses have been that estuaries are critical for calf rearing and that they provide protection from predators, warmer water temperatures, and an abundance of prey [5], but no study has examined the distribution and habitat use of mothers with calves. In the western and central Arctic, male and female belugas have shown differing distributions, with females with calves assumed to be spending more time in shallow waters [6]. In the Beaufort Sea, belugas have been shown to segregate by length (age), sex and reproductive status, where females with calves and small males select near shore open water habitat and larger males select ice covered offshore areas [9]. It is unknown, however, why these belugas segregate in the eastern Beaufort Sea but predation and foraging are two likely explanations [9]. In western Hudson Bay, foraging and predation may also explain beluga whale distribution.

Belugas have been observed feeding on capelin within estuaries [10, Westdal et al., unpublished data], they remain relatively close to shore throughout the summer season [5, 11] and females and calves appear to concentrate in the Churchill and Seal River estuaries in July perhaps due to the risk of killer whales predation [12]. The predation risk hypothesis suggests that in a slow reproducing species such as the beluga, behaviour that minimizes predation risk, in this case possible selection of shallow water habitat for females with calves, is favored [13]. Here our objectives were to determine if there is a difference in habitat use by adult belugas with calves compared with other age classes in the Seal and Churchill River areas from aerial photos taken in 2015 and further, to investigate differences in environmental characteristics that might explain differences in habitat use by age class.

## Methods

An analysis of the Fisheries and Oceans Canada (DFO) 2015 Western Hudson Bay beluga aerial survey was undertaken to look at belugas by age class and their locations within the study area (which is considered to cover the majority of the summer range of the WHB beluga population). The survey took place between August 6^th^ and 19^th^ of 2015 and followed methods and covered an area similar to that of Richard [14]. (See [11]) for a detailed description of the survey). A visual as well as photographic survey took place in high-density areas, including the three main estuaries in western Hudson Bay, allowing for individual identification of belugas. Here we analyzed photos from the Seal and Churchill river estuaries and surrounding area.

Although close in geographic proximity, the Churchill and the Seal estuaries do not have similar coastlines and bathymetry. The intertidal zone, mud flat and shallow/foul ground waters (less than 5 meters at lowest normal tide) extend more than 10 km off shore from the mouth of the Seal River Estuary, whereas deeper waters (≥10 m at lowest normal tide) are found just outside the mouth of the Churchill River Estuary (Figure 1). For this reason, results from estuaries were not compared to each other.

**Figure 1.**
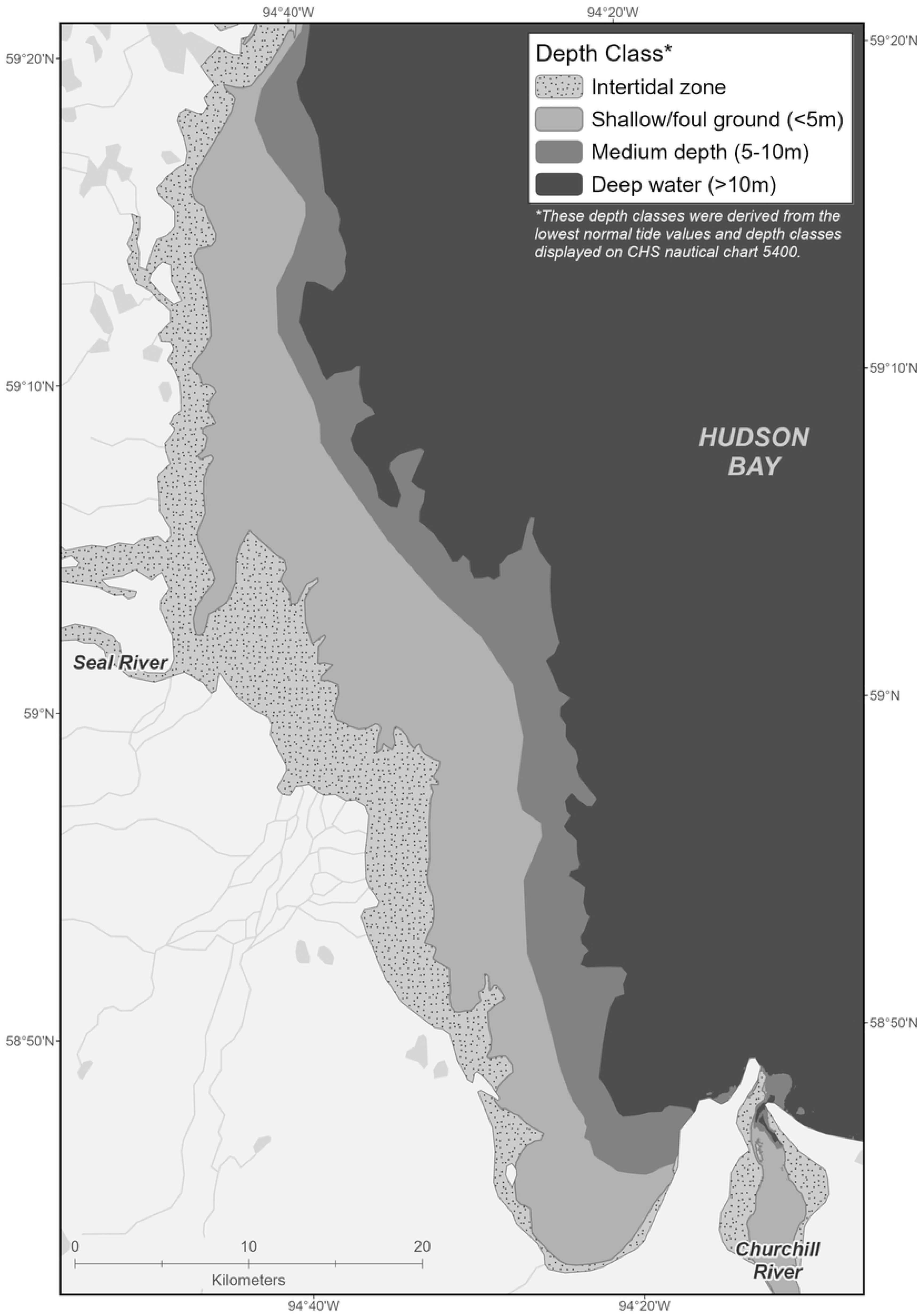

Photographs were taken using two Canon cameras (25 mm and 50 mm lenses) mounted on a twin otter flying at 2,000 feet over the Churchill and Seal River estuaries (complete photographic coverage) and offshore areas (transect coverage) (Figure 2). A review of photos from both cameras in the Churchill estuary determined that we were able to identify neonates in photos from the 25 mm lens and thus these photos were used in this analysis to provide more coverage than photos from the 50 mm lens. Individual photographs (n=3,661) covered a surface area of approximately 857 × 585 m with an estimated 20-40% overlap between photos.

**Figure 2.**
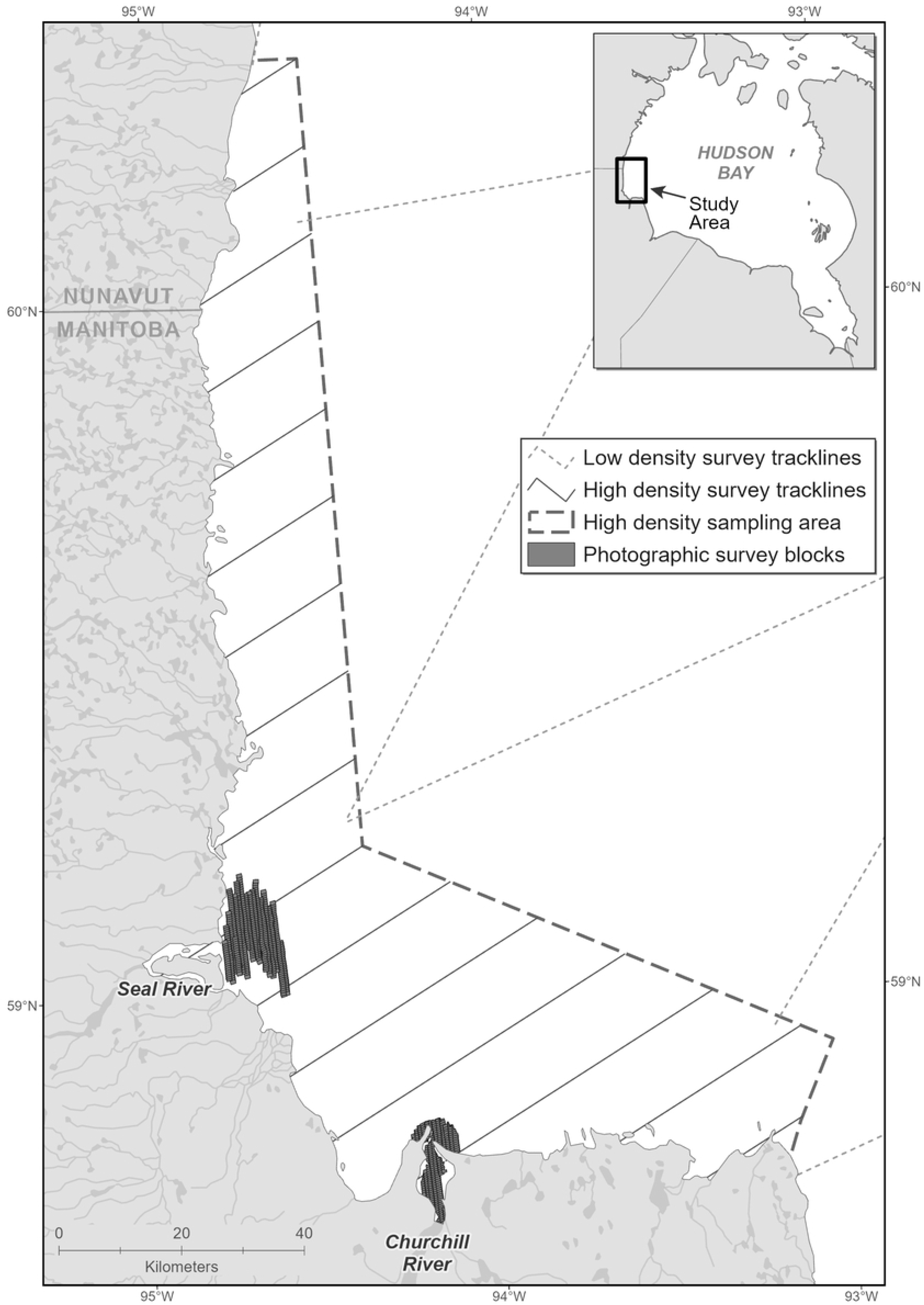

Photographs were scanned individually and belugas were identified and classified into three age categories (adult, juvenile, calf) visually. Calves and juveniles were distinguished from each other as well as adults using a variation on a dichotomous key developed to detect narwhal newborns in aerial photographs [15]. Calves were 33-50% the size of adult whales, less than two body lengths away from an adult whale and located adjacent to, above, on top or behind the nearest adult whale (ie. not ahead of). Juveniles were grey in coloration and 50 to 75% the size of adults and adults were the largest of the three age classes and white in color. Photos were georeferenced, GPS coordinates of identified calves, juveniles and adults were plotted in ArcGIS, and duplicates (from overlapping photos) were removed. A 500 m^2^ grid was laid on top of the survey area and each individual was assigned a cell. For each cell center, five variables were generated using geoprocessing tools in ArcGIS: average sea surface temperature [16], distance to shore and intertidal zone, distance to nearest river mouth (center of river mouth from headland to headland), and distance to minimum river plume extent (Atreya Basu, Unpublished Data, Center for Earth Observation Science). Each grid cell containing belugas was designated as belonging to either Seal (n=152), Churchill (n=60), or “outside”(n=11), depending on the location. Proportion of belugas identified by age class was calculated for the three identified areas and Fisher’s Exact Test compared age/sex class ratios between the two main estuaries, the Seal and Churchill River estuary. Beluga distribution by age class was examined in relation to distance to coastal habitat and bathymetry (predation risk hypothesis), sea surface temperature (thermal advantage), and extent of river plume (could relate to forge selection, predation risk and thermal advantage) using Kruskal-Wallis test run in JASP [17].

## Results

The total number of belugas counted, after removing duplicates from overlapping photos, was 13,538 (Table 1). Dense clumping of animals was common at the two estuaries, particularly near the Seal River Estuary (Figure 3).

**Table 1.** Total beluga whales identified (and proportion of total) in 2015 Western Hudson Bay survey (not including the Nelson River Estuary) by location and age class.

| Age Class | Location |  |  | Total number of animals |
| --- | --- | --- | --- | --- |
|  | Seal | Churchill | “Outside” |  |
| Adult | 11,115<br>(90.6%) | 1,031<br>(89.4%) | 106<br>(88.3%) | 12,252 |
| Juvenile | 629<br>(5.1%) | 52<br>(4.5%) | 6<br>(5%) | 687 |
| Calf | 521<br>(4.3%) | 70<br>(6.1%) | 8<br>(6.7%) | 599 |
| Total | 12,265 | 1,153 | 120 | 13,538 |

**Figure 3.**
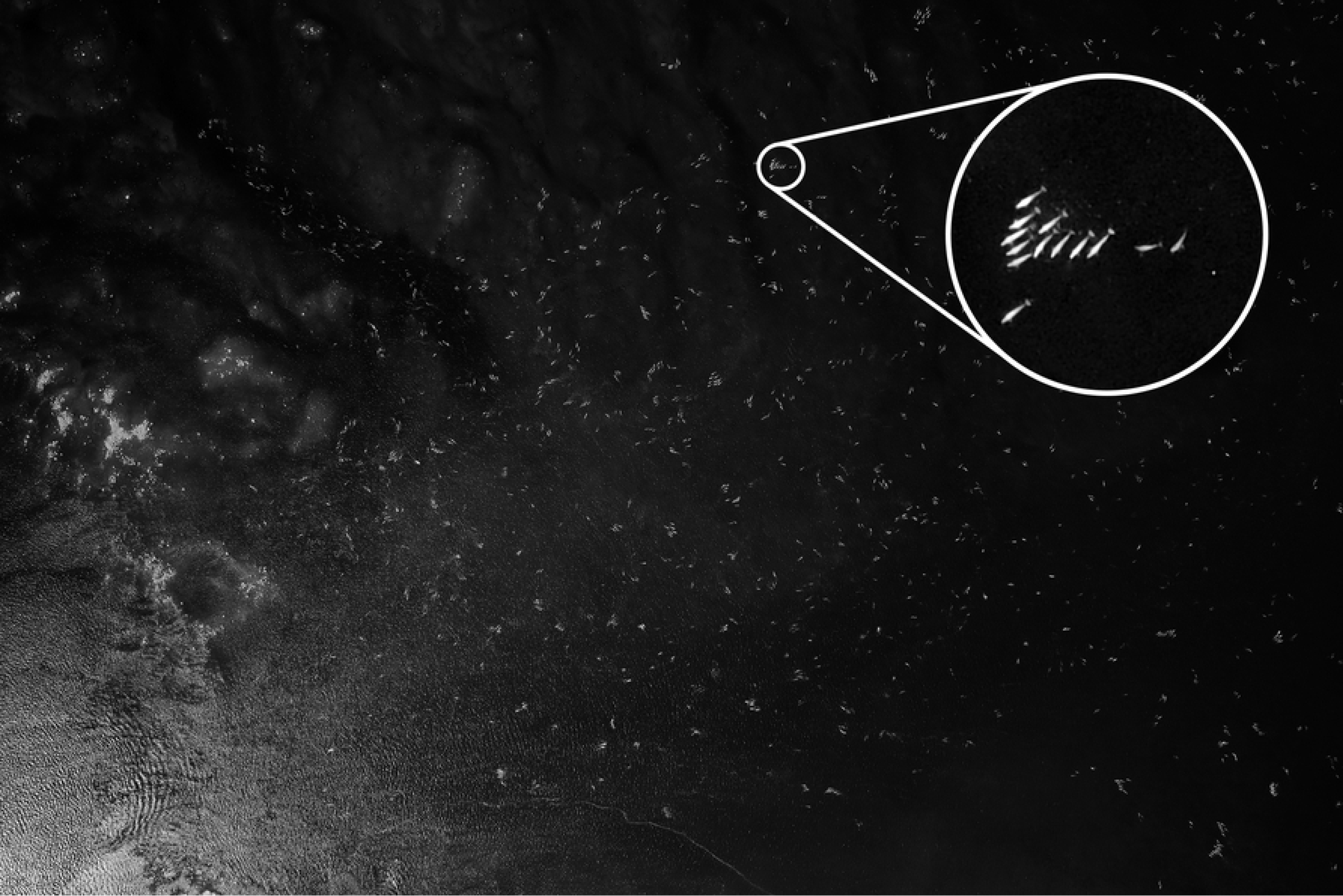

Beluga numbers were highest at the Seal Estuary (n=12,265), with few animals identified offshore (n=120) (Table 1). The Churchill and the Seal estuaries contained a similar proportion of beluga by age class. Calves made up 4.25% to 6.67% of the total beluga count in both estuaries, as well as the small number of belugas clumped between the two estuaries (Table 1).

Fisher’s Exact Test comparing the proportion of belugas by age class at the Seal and Churchill estuaries provided a p-value of 0.936, confirming similarity between estuaries.

All belugas observed were inside the river plume extent (Figure 4), with distances to the minimum plume extent (blue line) ranging from 4,810 m to 21,519m. Mean distance to intertidal, coast, river plume and river were calculated for each estuary by age class. Distances across age classes to geographical features were similar by estuary. Sea surface temperatures were similar in both locations and for all age classes (Table 2).

**Figure 4.**
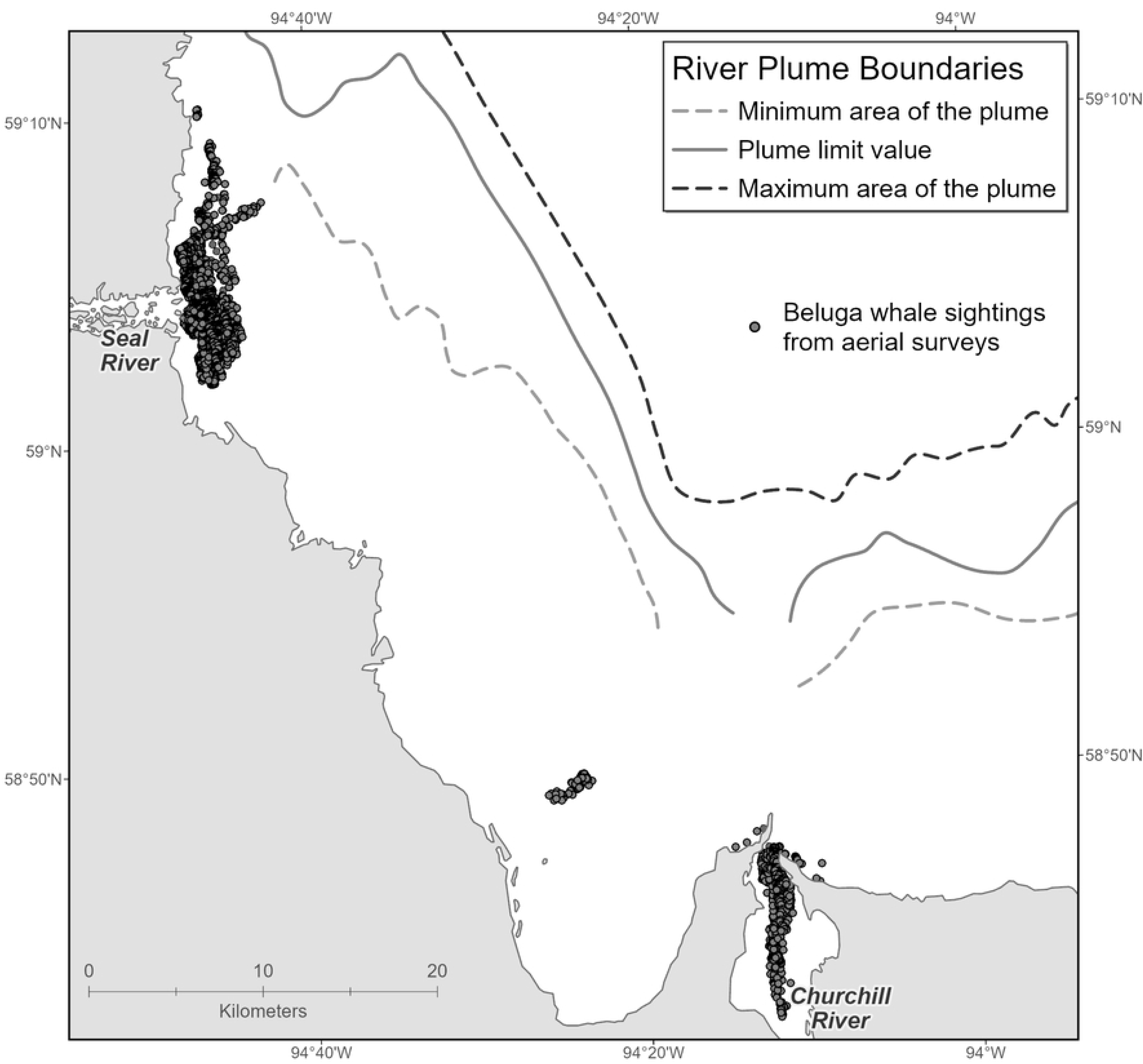

**Table 2.** Belugas by estuary and age class compared to environmental variables: median distance (25% and 75% quantile) to intertidal zone, distance to coastal, distance to river plume, distance to river and sea surface temperature.

| Age Class | Median distance (m) to intertidal zone |  | Median distance (m) to coast |  | Median distance (m) to river plume |  | Median distance (m) to river |  | Median SST (°C) |  |
| --- | --- | --- | --- | --- | --- | --- | --- | --- | --- | --- |
|  | Churchill | Seal | Churchill | Seal | Churchill | Seal | Churchill | Seal | Churchill | Seal |
| *Adult* | **600** | **0** | **825** | **1,204** | **14,342** | **12,298** | **3,114** | **2,631** | **12.38** | **12.14** |
|  | Q1: 316 | Q1: 0 | Q1: 447 | Q1: 640 | Q1: | Q1: | Q1: | Q1: | Q1: | Q1: |
|  | Q3: 806 | Q3: 0 | Q3: | Q3: | 13,354 | 11,241 | 2,110 | 2,102 | 12.34 | 12.12 |
|  | n=1031 | n=11,115 | 1,118 | 1,703 | Q3: | Q3: | Q3: | Q3: | Q3: | Q3: |
|  |  |  | n=1031 | n=11,115 | 16,224 | 12,971 | 4,105 | 3,289 | 12.44 | 12.15 |
|  |  |  |  |  | n=1031 | n=11,115 | n=1031 | n=11,115 | n=1031 | n=11,115 |
| *Juvenile* | **550** | **0** | **825** | **1,400** | **14,812** | **12,093** | **2,861.50** | **2,631** | **12.38** | **12.14** |
|  | Q1: | Q1: 0 | Q1: | Q1:707 | Q1: | Q1:11,37 | Q1: | Q1:2,102 | Q1: | Q1:12.12 |
|  | 408.25 | Q3: 0 | 435.25 | Q3:1,709 | 13,354 | 4 | 1,989.50 | Q3:3,228 | 12.34 | Q3:12.15 |
|  | Q3: 825 | n=629 | Q3: | n=629 | Q3:15,85 | Q3:12,77 | Q3: | n=629 | Q3: | n=629 |
|  | n=52 |  | 1,087.25 |  | 6.25 | 7 | 4,105 |  | 12.43 |  |
|  |  |  | n=52 |  | n=52 | n=629 | n=52 |  | n=52 |  |
| *Calf* | **500** | **0** | **825** | **1,476** | **14,821** | **12,093** | **2,617** | **2631** | **12.38** | **12.13** |
|  | Q1: 316 | Q1: 0 | Q1: 447 | Q1: 900 | Q1: | Q1:11,18 | Q1: | Q1: 2138 | Q1: | Q1:12.12 |
|  | Q3: 806 | Q3: 0 | Q3: | Q3: 1900 | 13,354 | 0 | 1,668.25 | Q3: 3289 | 12.34 | Q3:12.15 |
|  | n=70 | n=521 | 1107.75 | n=521 |  |  |  | n=521 |  | n=521 |
|  |  |  | n=70 |  | Q3:16,09 | Q3:12,77 | Q3: |  | Q3: |  |
|  |  |  |  |  | 2.75 | 7 | 4,371.75 |  | 12.45 |  |
|  |  |  |  |  | n=70 | n=521 | n=70 |  | n=70 |  |

Box plots (Figure 5) showed the similarity of means and overlapping standard deviations for age classes of each environmental variable.

**Figure 5.**
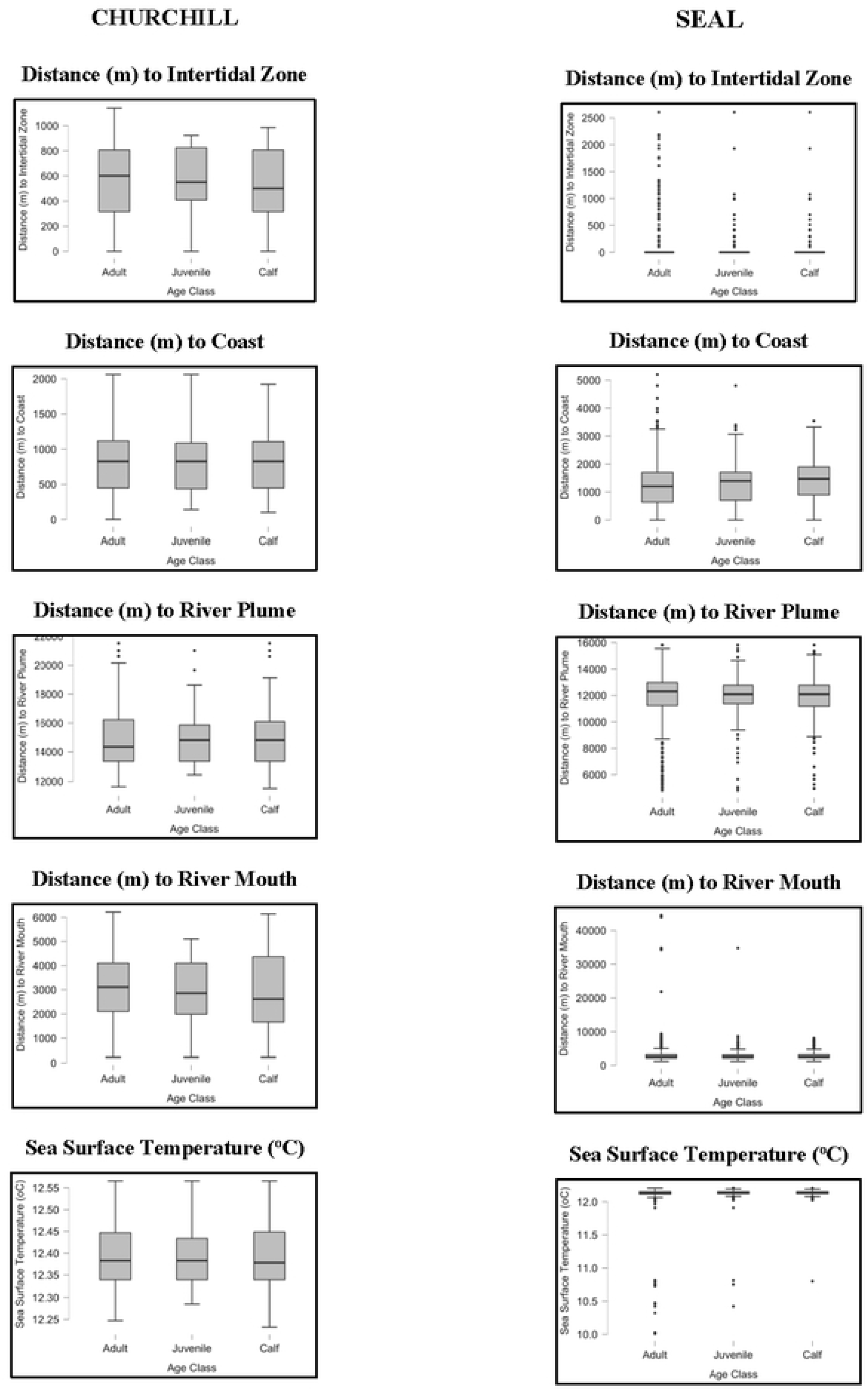

Kruskal-Wallis test showed that there was only one significant variation between age classes and environmental variables, distance to coast near the Seal River (Table 3).

**Table 3.** Kruskal-Wallis Test comparing differences between environmental co-variates and age class of belugas near the Churchill and Seal River Estuaries.

|  |  | **Statistic** | **df** | **p** |
| --- | --- | --- | --- | --- |
| **Intertidal** | Churchill | 0.367 | 2 | 0.83 |
|  | Seal | 0.867 | 2 | 0.65 |
| **Distance to Coast** | Churchill | 0.136 | 2 | 0.93 |
|  | Seal | 23.311 | 2 | <0.001* |
| **Distance to Plume** | Churchill | 0.417 | 2 | 0.81 |
|  | Seal | 3.623 | 2 | 0.16 |
| **Distance to River** | Churchill | 0.548 | 2 | 0.76 |
|  | Seal | 2.101 | 2 | 0.35 |
| **Sea Surface Temperature** | Churchill | 0.333 | 2 | 0.85 |
|  | Seal | 1.423 | 2 | 0.49 |

Distance to coast by age class for Seal River estuary beluga did not differ among age class, however the distribution was less uniform (Figure 6).

**Figure 6.**
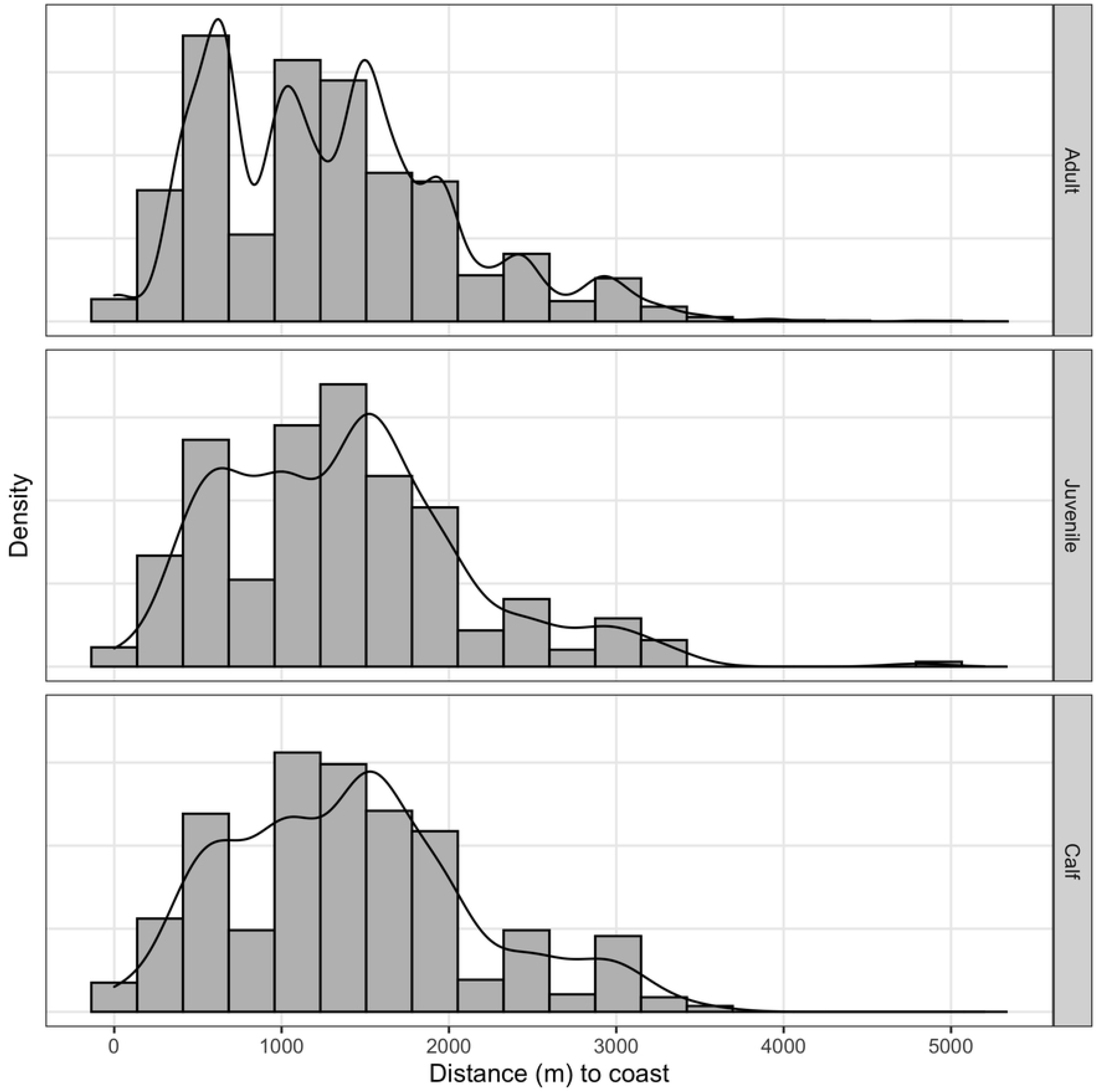

## Discussion

The Western Hudson Bay beluga population gathers in dense groups in the summer season. This analysis considered environmental factors as possible drivers of distribution and showed that not any one of these factors was significant, and adults with calves did not select separate habitat from juvenile and adults. All beluga groups were observed well inside the river plume extent, sea surface temperatures were similar across the study area, and distances to geographical features were similar across age classes by estuary. Similar to our findings, Kelley and Ferguson [18] found that male and female belugas (from Eastern and Western Hudson Bay beluga populations), in and out of mating season, appeared to be foraging similarly and perhaps occupying the same habitat; however Barber et al. [6] (Eastern Beaufort Sea and Eastern High Arctic-Baffin Bay) and Loseto et al. [9] (Eastern Beaufort Sea) both found segregation of belugas by age or sex.

Cetaceans are known to segregate by sex and age. Adult male sperm whales (*Physeter microcephalus*) separate from females, calves and juveniles [19], Hector’s dolphin (*Cephalorhynchus hectori*) groups segregate by sex [20], and bowhead whales segregate geographically by sex [21]. Periodic segregation has also been recorded in humpback whales during migration and in killer whales when foraging [22]. Narwhal, the closest relative of beluga, appear to travel in groups of mixed sex and age class [23], but diet may differ by sex [18].

Proportion of belugas by age classes were similar between the Churchill and Seal River estuaries, and the cluster of animals between the estuaries in Button Bay, suggesting that one location is not preferred over another for calf rearing in the summer season. A range of 4.3%-6.1% of visible whales were calves across the three identified regions, indicating that location is not a driver of calf abundance. Exhibiting high site fidelity, belugas are thought to return to the same locations seasonally each year [24]. In this region, beluga with markings have been resighted year to year (S. Peterson, personal communication, K. Westdal, personal observations) and recent Western Hudson Bay beluga population estimates are also similar (57,300 (95% CI: 37,700-87,100) [14], 54,473 (95% CI: 44,988-65,957) [11].

Distance to key environmental features did not differ among age classes suggesting a mixed pattern of distribution. Identified belugas were inside the river plume, sea surface temperature was similar across the study area, and distances to land and or bathymetric locations were not significant across age classes. Distance to coast was a significant factor between age classes for the Seal River, however bathymetry in this region could explain the variation with mud and tidal flats extending greater than 10 km offshore non-uniformly. Beluga whales remained in the same habitat type, intertidal and shallow or foul ground (less than 3 meters at average low tide) despite the variation. This bathymetry, which is different from that near the Churchill Estuary, did not allow for a comparison from one estuary to another.

The highly social nature of belugas [25], like other cetaceans, may make it difficult to explain abundance and distribution in relation to environmental characteristics [26]. Marine mammals are known to congregative in large aggregations that are not clearly tied to habitat [26]. Beluga may be gathering in large groups in the summer season for social rather than environmental reasons. Rendell and Whitehead [27] suggest that particular locations may be selected seasonally as a result of cultural traditions. Beluga migration routes and seasonal habitat association appear to be passed down through females to offspring in belugas [28].

Uncertainties remain however based on environmental data available and nature of the beluga photographic survey data. Prey distribution and salinity were not included as environmental variables due to a lack of data availability. We were also using data from one survey in one year, providing a snap shot in time. Anderson et al. [29] for example found that beluga groups with calves in Cunningham Inlet were found closer to shore on cloudier days (possible terrestrial anti-predation behaviour). And Smith et al. [5] found that beluga distribution from the Nelson river mouth varied depending on fresh water outflow volumes from the river by year.

This study offers evidence to suggest that age segregation is not occurring in the Western Hudson Bay beluga population at this time of year. Here adults with calves did not choose shallower waters more frequently than other age classes, or congregate at the river mouths. The reproductive risk-predation risk hypothesis suggests that belugas with calves would choose shallower waters [30] where killer whales would be excluded (although belugas would still have access to this habitat). Beluga distribution also did not extend out into the river plume (where further feeding opportunities may be possible). Forage-selection hypothesis predicts a difference in distribution by sex where females select for quality vs males selecting habitat based on forage quantity [31]. Temperatures were fairly consistent throughout the entire survey area, suggesting that thermal advantage, or the selection of warmer waters isn’t part of site selection and not necessary for beluga neonates as suggested by Doidge et al. [32]. Selection of shallow waters may play a role in habitat selection, but unlikely to be the significant determinant (as suggested by Smith et al. [5]) based on the distribution of belugas and lack of differentiation seen amongst age classes.

Future studies could include focal follow studies of females and calves, remotely deployed satellite telemetry tags on females with calves, as well as longer time frame video monitoring to look at group structure over time. Further understanding is needed of social behaviour and habitat use over the full summer season, as well as age class distribution in the third major estuary utilized by beluga in the region (Nelson River estuary). Our results provide new information on beluga habitat use for conservation and management planning in the light of provincial government interest in protection of upstream habitat and Federal government interest in coastal marine protection in Western Hudson Bay.

## Acknowledgements

Survey data was collected and analysis was funded by Fisheries and Oceans Canada. Thank you to Leah Braithwaite and Chris Paetkau for their roles in review of survey photographs and special thanks to Heather Grant for photo editing, Olivia Mussels at Oceans North for early GIS assistance and Claire Hornby at Fisheries and Oceans Canada for last minute R support.

## References

1. Ruckstuhl KE, Neuhaus P. Sexual segregation in vertebrates: ecology of the two sexes. Cambridge University Press. 2005. doi: 10.1017/CBO9780511525629

2. Cosens SE, Blouw A. Size and age class segregation of bowhead whales summering in northern Foxe Basin: A photogrammetric analysis. Mar Mamm Sci. 2006;19(2): 284–296.

3. Hamilton PK, Cooper LA. Changes in North Atlantic right whale (Eubalaena glacialis) cow-calf association times and use of the calving ground: 1993-2005. Mar. Mamm. Sci. 2010;26(4): 896–916.

4. Urban RJ, Rojas-Bracho L, Perez-Cortes H, Gomez-Gallardo A, Swartz SL, Ludwig S, Brownell Jr RL. A review of gray whales (Eschrichtius robustus) on their wintering grounds in Mexican waters. J. Cetacean Res. Manage. 2003;5(3):281–295.

5. Smith AJ, Higdon JW, Richard P, Orr J, Bernhardt W, Ferguson SH. Beluga whale summer habitat associations in the Nelson River estuary, Western Hudson Bay, Canada. PLoS ONE. 2017;12(8): doi: 10.1371/journal.pone.0181045

6. Barber, DG, Saczuk, E, Richard, PR. Examination of beluga-habitat relationships through the use of telemetry and a Geographic Information System. Arctic. 2001;54:305–316.

7. Colbeck GJ, Duchesne P, Postma LD, Lesage V, Hammill MO, Turgeon J. Groups of related belugas (Delphinapterus leucas) travel together during their seasonal migrations in and around Hudson Bay. Proc R Soc. 2013;280. doi: 10.1098/rspb.2012.2552

8. Luque SP, Ferguson SH. Age structure, growth, mortality, and density of belugas (Delphinapterus leucas) in the Canadian Arctic: responses to environment? Polar Biology. 2010;33(2):163–178.

9. Loseto LL, Richard P, Stern GA, Orr J, Ferguson SH. Segregation of Beaufort Sea beluga whales during the open water season. Can. J. Zool. 2006;84:1743–1751.

10. Watts PD, Draper BA. Note on the behaviour of beluga whales feeding on capelin. Arctic and Alpine Research. 1986;18(4):439.

11. Matthews CJD, Watt CA, Asselin NC, Dunn JB, Young BG, Montsion LM, Westdal KH, Hall PA, Orr JR, Ferguson SH, Marcoux M. Estimated abundance of the Western Hudson Bay beluga stock from the 2015 visual and photographic aerial survey. DFO Can. Sci. Advis. Sec. Res. Doc. 2017/061. v + 20 p.

12. Westdal KH, Davies J, MacPherson A, Orr J, Ferguson SH. Behavioural changes in Belugas (Delphinapterus leucas) during a Killer Whale (Orcinus orca) attack in southwest Hudson Bay. Canadian Field-Naturalist. 2016;130(4):315–319.

13. Michaud R. Sociality and ecology of the odontocetes. In Sexual segregation in vertebrates: Ecology of the two sexes. Edited by K.E. Ruckstuhl and P. Neuhaus.Cambridge University Press, 2005:303–326.

14. Richard PR. An estimate of the western Hudson Bay beluga population size in 2004. Canadian Science Advisory Secretariat Research Document 2005/017.

15. Charry B, Marcoux M, Humphries MH. Aerial photographic identification of narwhal (Monodon monoceros) newborns and their spatial proximity to the nearest adult female. Arctic Science. 2018;4:513–524

16. JPL MUR MEaSUREs Project. 2010. GHRSST Level 4 G1SST Global Foundation Sea Surface Temperature Analysis. Ver. 1. PO.DAAC, CA, USA. Dataset accessed [2020-02-03] at https://doi.org/10.5067/GHGMR-4FJ04

17. JASP Team. JASP (Version 0.14.1) 2020 [Computer software]

18. Kelly TC, Ferguson SH. Sexual segregation in two closely related species: Beluga whales (Delphinapterus leucas) and narwhal (Monodon monoceros). PhD Thesis, The University of Manitoba. 2014. Available from: https://mspace.lib.umanitoba.ca/bitstream/handle/1993/23548/kelley_tritsya.pdf?sequence=1

19. Christal J, Whitehead H. Social affiliations within sperm whale (Physeter microcephalus) group. Ethology. 2001;107:323–240.

20. Webster TA, Dawson SM, Slooten E. Evidence of sex segregation in Hector’s dolphin (Cephalorhynchus hectori). Aquatic Mammals. 2009;35(2):212–219.

21. Heide-Jorgensen MP, Laidre KL, Wiig O, Postma L, Dueck L, Bachmann L. Large-scale sexual segregation of bowhead whales. Endangered Species Research. 2010;13:73–78.

22. Beerman A, Ashe E, Preedy K, Williams R. Sexual segregation when foraging in an extremely social killer whale population. Behavioral ecology and sociobiology. 2016;70(1):189–198.

23. Marcoux M, Auger-Methe M, Humphries MM. Encounter frequencies and grouping patterns of narwhals in Koluktoo Bay, Baffin Island. Polar Biology. 2009;32:1705–1716.

24. COSEWIC. COSEWIC assessment and status report on the Beluga Whale Delphinapterus leucas, Eastern High Arctic - Baffin Bay population, Cumberland Sound population, Ungava Bay population, Western Hudson Bay population, Eastern Hudson Bay population and James Bay population in Canada. Committee on the Status of Endangered Wildlife in Canada. 2020 In Press. Ottawa. xxxv + 84 pp.

25. Kingsley M, Gosselin S, Sleno G. Movements and Dive Behaviour of Belugas in Northern Quebec. Arctic. 2001;54(3):262–275.

26. Whitehead H, Parijs SV. Studying marine mammal social systems. In: Boyd I, Bowen D, Iverson S, editors. Marine mammal ecology and conservation: a handbook of techniques. Oxford University Press, Oxford, U.K.; 2010. pp. 263–282.

27. Rendell L, Whitehead H. Culture in whales and dolphins, Behav. Brain Sci, 2001;24:309–382.

28. O’Corry-Crowe GM, Suydam RS, Rosenberg A, Frost KJ, Dizon AE. Phylogeography, population structure and dispersal patterns of the beluga whales Delphinapterus leucas in the western Nearctic revealed by mitochondrial DNA. Molecular Ecology. 1997;6:955–970.

29. Anderson PA, Poe RB, Thompson LA, Webber N, Romano TA. 2017. Behavioral responses of beluga whales (Delphinapterus leucas) to environmental variation in an Arctic estuary. Behav. Process. 2017;145:48–59.

30. Grignolio S, Rossi I, Bassano B, Apollonio M. Predation risk as a factor affecting sexual segregation in Alpine Ibex. Journal of Mammalogy. 2007;88(6):1488–1497.doi: 10.1644/06-MAMM-A-351R.1

31. Ruckstuhl KW, Neuhaus P. Sexual segregation in ungulates: a comparative test of three hypotheses. Biol Rev Camb Philos Soc. 2002;77(1):77–96.doi: 10.1017/s1464793101005814.

32. Doidge DW. Integumentary heat loss and blubber distribution in the beluga, Delphinapterus leucas, with comparisons to the narwhal, Monodon monoceros. In: Smith TG, St. Aubin DG, Geraci JR, editors. Advances in research on the beluga whale, Delphinapterus leucas. Can Bull Fish Aquat Sci; 1990. pp. 129–140.

